# Evaluation of cytotoxic, antiviral effect and mutagenic potential of a micronutrient combination in vitro cell culture

**DOI:** 10.1101/2020.06.18.160333

**Authors:** Nazish Matti, Muhammad Ashraf, Muhammad Adil Rasheed, Imran Altaf, Isabel Carvalho, Muhammad Faisal Nadeem

## Abstract

**Background:** Micronutrients are essential for the body to produce enzymes, hormones and other substances essential for proper growth and development. Iodine, potassium iodide and ascorbic acid are ones of the most important in global public health terms. However, their lack represents a major threat to the human and animal health.

**Methodology:** Antiviral, cytotoxic and mutagenic activity of commercially available micronutrient combination consisting of iodine, ascorbic acid, potassium iodide and excipients was evaluated. Commercial preparation of micronutrient combination was compared with self-prepared micronutrient preparation. MTT assay was used to evaluate the cytotoxicity of all the preparation. Confluent monolayer of Chicken Embryo Fibroblasts grown in 96-well cell culture plates were treated with ten concentrations of each preparation in triplicate manner and was used to determine the viability of the cells and cell survival percentage. Antiviral efficacy was determined against influenza virus H9N1 strain by virus infection and subsequent cell viability assay. Furthermore, mutagenicity was measured by bacterial reverse mutation analysis by Ames test using two strains of *Salmonella typhimurium* TA100 and TA 98 with and without S9. After appropriate incubation, number of revertant colonies per plate were counted in triplicate manner and MF was calculated.

**Results:** Our combination showed cytotoxicity at doses higher than 11.71ug/ml while showed significant antiviral efficacy at concentrations of 11.71ug/ml, 5.86ug/ml, 2.93ug/ml and 1.4ug/ml which faded away at lower dilutions. Commercially available preparation was found to be non-mutagenic in our experiment.

**Conclusions:** Combination consisting of iodine, ascorbic acid, potassium iodide is effective in the treatment of viral infections.

## Introduction

Chronic viral hepatitis infection, characterized by liver inflammation, may lead to serious complications. One of the major causative agents is Hepatitis C Virus (HCV) which is the most common blood borne chronic infection in humans (Maghlaoui et al. 2012). Around the globe, approximately 130-170 million people are harboring the virus in them chronically and are destined to face the consequent complications while 350,000 die annually due to HCV infection. Along with high mortality, chronic HCV infection also carry huge financial burden due to its chronic nature and consequent complications. Research reports suggest the Hepatitis C virus incidence at 4.95% of the population implying that approximately 10 million people are infected with the virus while equivalent number is thought to be undiagnosed at this point of time (Ali et al, 2009).

Current standard treatment strategy for HCV involves the combination of pegylated interferon (PEG-IFN) either alpha-2 or beta 2 and Ribavirin (RBV) for 6-12 months. New pharmacotherapeutic agents, vaccines and biological agents are being discovered while goal for next decade has been set to interferon-free pharmacotherapy (Dore, 2012). Oral HCV NS3 protease inhibitors boceprevir and telaprevir (approved by FDA for use in human) when combined with current interferon and ribavirin therapy are more effective but increasing cost and toxicity pose a significant challenge in its widespread use (Assis and Lim, 2012).

Micronutrients are essential for the body to produce enzymes, hormones and other substances essential for proper growth and development (Aragony, 2013). Their properties, particularly the binding to large molecules, are far from specific, an observation which is reflected in the very wide range of diseases in which trace elements are employed. Micronutrient deficiencies have been reported in many diseased states (Harika, 2017; Shivakoti, 2016). These micronutrients (vitamins and trace elements) can alter biochemical processes. Guidelines for 13 essential vitamins and 10 essential trace elements have already been recognized while rest are being established in healthy individuals while diseased person may require more of these (Al-Sonboli et al. 2009).

Micronutrients, mainly believed to be essential for normal functioning of the body could possess potential beyond mere daily requirements. One of the such micronutrient combinations intended to be used for therapeutic purposes Commercial iodine complex (MTI medical)® which consist of iodine, potassium iodide and ascorbic acid while the main ingredient for its activity is believed to be polyiodides created by combination. When combined with current standard regime of interferon and Ribavirin, it is claimed to increase antiviral efficacy and decrease duration of treatment. This introduction demands tenacious proof of useful antiviral activity if any, at therapeutic doses to validate the claims of manufacturer. Although, the results merely indicating useful antiviral activity alone might not be enough to warrant the use of Commercial iodine complex (MTI medical) as antiviral agent. An open labelled, active controlled clinical trial was published regarding use of commercial ioding complex named is RENESSANS in chronic hepatitis C patients and it showed highly promising results. In the light of these results, it seemed interesting to investigate seaweed iodine complex which could demonstrate selective toxicity against viruses (Hassan, 2015). In our laboratory, we have already developed avian influenza virus model which could be used for investigational purpose.

**Vitamin C** (ascorbic acid) has long been known for its antiviral effects. It has been implied in nutritional therapies for viral infections at therapeutic diseases. Ascorbic acid was also shown to overcome the phenotypic expressions in cells also known as cytopathogenic activity of viruses in multiple cells (Lima et al. 2010). Vitamin C, at therapeutic doses, causes significant decrease in ability of avian tumor viruses to multiply in cultured cells. Ascorbic acid has long been known as cytoprotective, antioxidant agent. It has been utilized safely in great quantities. High doses taken orally, and IV injections have produced concentrations of up to 250uM/L which does not appear cytotoxic in many studies (Levine et al, 2011).

**Iodine** is considered an excellent antimicrobial and specifically antiviral action of the element commonly used in nutrition. Cell culture studies also show that higher concentrations of iodine have exceptional antiviral activities. Recently, the potential of iodine against Human Immunodeficiency Virus (HIV) have been tested due to its powerful antiviral activity. Encouraging results were shown in 9 terms of HIV suppression, reduced infectivity and oral bioavailability of the iodine (Mamo & Naissides, 2005). It highlighted some important aspects of mechanism of action of iodine as antiviral agent (Sriwilaijaroen et al. 2009). Excessive iodine can be obtained from dietary supplements, natural foods, topical antiseptics and contrast media. This increased amount may result in thyroiditis, sensitivity reactions, hypo or hyperthyroidism or idiosyncratic reactions in few (Patrick, 2008). Children with high thyroid activity and immature follicle cells are more at risk compared to adult population (Leger & Carel, 2013).

The current project is designed to evaluate some of pharmacological properties of Commercial iodine complex which might form the basis of further testing and may direct the future course. First of all, its verifiable antiviral activity needs to be assessed while the second part is concerned with the safety of therapeutic concentrations. Importance of safety profile is also highlighted by the fact that previous assessment of iodinated compounds has questioned systemic use of these compounds due to thyroid toxicity and other side effects. On the other hand, ascorbic acid is quite safe even at higher doses.

The objectives of this study are to evaluate cytotoxicity of this combination by in vitro cell culture using MTT assay, characterize the antiviral activity of branded micronutrient combination Commercial iodine complex and to evaluate the mutagenic potential of this drug by Ames test.

## Materials and methods

### 3.1 Experimental design

The research was designed to determine the antiviral activity of Commercial iodine complex (MTI Commercial micronutrient combination), self-prepared similar micronutrient combination (UVAS Commercial iodine complex) locally prepared, individual ingredients of micronutrient combinations i.e. ascorbic acid and iodine notably along with positive or negative controls and antiviral standards. As standard antiviral drugs like Ribavirin, Oseltamivir and Amantadine were also tested. For Ames test two-fold 10 dilutions were evaluated for Commercial iodine complex and UVAS-iodine complex while individual ingredients were measured up to 6 doubling dilutions. Standard antiviral drugs were tested for 14 doubling dilutions. While in cytotoxicity and antiviral assays, 10 two-fold dilutions of each ingredient and standards were utilized by using chicken primary fibroblast cell line against H9 influenza virus. Different concentrations were made in cell culture media at the time of MTT assay by using stock solution and same dilutions were made in normal saline for antiviral activity. In cytotoxicity assay monolayer of primary fibroblast cells grown in 96-well cell culture plate were treated with each concentration. In this assay primary fibroblast cells along with cell culture media were kept as positive control however primary fibroblast cells, DMSO 10% and cell culture media were taken as negative control respectively. Viability of cells was checked by MTT calorimetric assay (Twentyman and Luscombe 1987).

### 3.2 AMES salmonella/microsome mutagenicity assay

Mutant strains of *Salmonella* typhimurium TA 100 and TA 98 were supplied by EBPI Canada in lyophilized form. Freeze dried *Salmonella typhimurium* strains were poured with 1ml of autoclaved nutrient broth in Biological Safety Cabinet II under aseptic conditions. After rehydration of almost 2 minutes each culture was dispensed to 5ml nutrient broth in screw cap glass tube. This culture was incubated at 37°C in shaking incubator shaken at 70 rpm for 24 to 36 hours. After that, the growth of *Salmonella typhimurium* TA 100 and TA 98 was visually checked. The growth of both strains was confirmed by streaking the cultures separately on nutrient agar plates which were previously incubated at 37°C to check the sterility. Then the streaked nutrient agar plates were placed in incubator at 37°C overnight.

### 3.3 Genetic Analysis

Individual cultures of TA 100 and TA 98 were subjected to genetic analysis for Histidine dependence, Biotin dependence, Histidine and Biotin dependence, rfa marker and pKM101 plasmid. To check that either bacteria are dependent on Biotinand/or histadine for their growth or not, GM agar plates containing excess of Biotin/Histadine (8ml of 0.01% w/v) were prepared as explained before. Each plate was streaked with a loopful of TA 100 and TA 98 individually under aseptic conditions. Then plates were incubated overnight at 37°C.

PKM101 Plasmid *Salmonella* TA 100 and TA 98 cultures were streaked on GM plates containing excess of Histidine and Biotin. A filter paper disc already soaked with 10μg/ml of Ampicillin was embedded on the streak of *Salmonella* to check either the TA 100 or TA 98 strain contained pKM101 plasmid or not and whether the mutant strains were resistant to Ampicillin as compared to the wild type of *Salmonella*.

### 3.4 Experimental Procedure

In this project Standard Plate Incorporation Method and Pre-incubation Assay were used to check mutagenicity of the test chemical. Pre-incubation Assay was applied when the chemical was assayed with S9-mix. Whereas Standard Plate Incorporation Assay was utilized when the test doses of chemical were checked without the addition of S9-mix.

First step in both the procedures is same. On the other hand, the second step was different in both Standard Plate in Corporation Assay & Pre-incubation Assay. After 48-hour incubation, colonies of revertant were counted by placing the plates on the Digital Colony Counter and results were expressed as Number of Revertant Colonies per plate. Ethidium Bromide and Nazide in the concentration of 5μg/plate was used as standard positive concentration to cause mutagenicity in both strains of *Salmonella typhimurium*.

*Salmonella* culture, media, Histidine and Biotin were added in the negative control plate. There was no drug chemical present in negative control plates (Mortelmans and Zeiger 2000).

### 3.5 Cytotoxicity assay

In cytotoxicity assay, confluent monolayers of Chicken Embryo Fibroblast (CEF) cells grown in 96-well cell culture plates were treated with each test concentration. In the assay, CEF cells along with cell culture media were kept as positive control, however CEF cells, DMSO (10%) and cell culture media were taken as negative control respectively. Viability of CEF cells was checked by using MTT [(3-(4,5dimethylthiazole–2 yl)-2,5 diphenyltetrazolium bromide)] calorimetric assay (Twentyman et al. 1987).

Cell culture media was prepared by adding 1.2g of the powdered M-199 in 100 ml of double distilled water along with antibiotics and fetal bovine serum (10% serum for growth media while 1% fetal bovine serum for maintenance media) was added to it. The media was then filtrated by using a negative pressure filtration assembly, in a safety cabinet.

#### 3.5.1 Preparation Chicken Embryo Fibroblast Cell Line

Nine-day embryos isolated from healthy non-infected chicken eggs (obtained from VRI, UVAS). Eggs were rinsed three times with PBS (pH 7.4) and then chopped into 1-mm3 pieces. Then these cells were placed in tissue cultivation flasks containing (minimal essential medium) +10% (v/v) fetal bovine serum along with Penicillin (1000 u/mL) and Streptomycin (1000 μg/ml) in a 75 cm2 flask in an incubator at 37°C with 5% CO2 in air (Freshney, 2010). Primary cells were subculture further into flasks when 80–90% confluent in the ratio of 1:2 or 1:3. All experimental cells were treated when they were ∼80% confluent.

#### 3.5.2 Quantification of CEF cells

To quantify cells a well cleaned hemocytometer was taken, and a cover slip was affixed on it. A drop of cell suspension was mixed with a drop of 0.4% Trypan blue, then loaded in the counting chamber of hemocytometer and was kept untouched for 1 to 2 minutes. The hemocytometer was placed on microscope. Both stained and unstained cells were counted. Stained cells represented dead cells and unstained cells represented viable cells (Freshney, 2010). Following formula was used to calculate % viable cells.

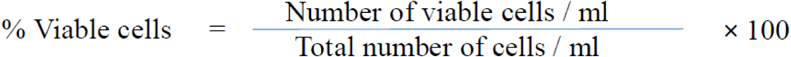

### 3.6 MTT assay for cytotoxicity

Two plates with 80-90% confluent monolayer of CEF cells were used for cytotoxicity assay and labeled as “cytotoxicity assay plate A”, and “cytotoxic assay plate B”. Cell culture media was removed from each well of the plates. The wells were washed with phosphate buffer saline solution (PBS). 100ul of fresh maintenance media and then 100ul of the extract was added to separate wells. Triplicate wells were used for cytotoxicity analysis of test concentration. Each plate was then covered with its lid and incubated at 370C and 5% CO2. After an incubation of 5-6 days, media was removed from each well of the plates. Fresh maintenance media was added after rinsing the monolayers with PBS. Then 100μl of 0.5% MTT solution was added to the test and control wells of cytotoxic assay plates and incubated at 370C for 3-4 hours. Cell culture media and MTT was then removed from each well and 100ul of DMSO (5%) was added to each well of the plates to dissolve the formazan crystals. The plates were kept in an incubator at 370C for 2 hours. Multi well ELISA reader at 570nm was used to measure the optical density (OD) values of each well of the plate (Twentyman and Luscombe, 1987). In this study, we used a Positive Control (PBS) and a negative one (10% DMSO). The Cell Survival Percentage (CSP) was obtain using the following:

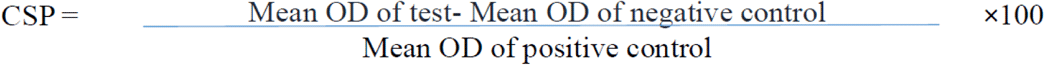

### 3.7 Antiviral Assay

Chicken Embryo Fibroblast Cells (CEF) were grown in Dulbecco’s Modified Eagle’s Medium (DMEM) (Mediatech Cellgro, USA), supplemented with 10% Fetal Bovine Serum (FBS) (PAA, Austria) and 1% Pen/Strep (Mediatech Cellgro, USA) at 37°C in a humidified incubator. The media was changed two to three times per week. The influenza virus strain, (H9N1), was obtained from American Type Culture Collection (ATCC) courtesy from WTO laboratories UVAS. It was propagated in MDCK cells in the presence of 1μg/ml of Trypsin_TPCK (Tosylamide Phenylethyl Chloromethyl Keton-treated Trypsin) (Sigma, USA) to create the working stock. During antiviral evaluations, media supplemented with FBS was sucked out and the cell was washed with PBS and then it was treated as needed. Then media supplemented with Trypsin_TPCK was added.

For the virus characterization, RNA extraction methodology was applied according with the manufacture. In order to quantify the nucleic acid concentration and level of purity, the absorbance readings were taken at 260 and 280nm (Spectrophotometer 132 ND-100, Nanodrop).

The presence of different genes was tested by PCR assays using PCR Thermal Cycler, WTO QOL UVAS, followed by electrophoresis. One set of primer was designed from segment 4 hemagglutinin (HA) gene from Avain Influenza. The primer was designed by using GenBank accession number JQ905259. The primer (Forward primer 5’ATC GGC TGT TAA TGG AAT GTG 3’) and (Reverse primer 5’TGG GCG TCT TGA ATA GGG TAA 3’) was used to amplify the partial HA gene (221 bp). The primer designing was done by using Primer3 software (http://frodo.wi.mit.edu/).

#### 3.7.1 HA titre for antiviral activity

Hemagglutination test was carried out to quantify the amount of H9N1 influenza virus in the suspension and to evaluate the antiviral activity of micronutrient combinations, individual ingredients and standard antiviral drugs. This was done by carrying out two-fold dilution of the viral suspension in a micro-well plate and then testing the end point, which is the lowest concentration of virus where there was hemagglutination. This result can then be used to determine the amount of haemagglutinin in the suspension and is expressed as a HA titre.

A total of 5ml of blood was taken from wing vein of an adult broiler chicken bird in a test tube containing EDTA as an anticoagulant. Chicken blood sample was centrifuged at 1500 rpm for 5 minutes. Then plasma was separated by sterile plastic dropper.

#### 3.7.2 Measurement of Antiviral activity by Cytotoxicity assay

CEF cells were incubated in 96-well micro-plate for 24 hours at 37°C. Two-fold serial dilutions of compounds were added to the semi-confluent cells in triplicates and incubated at different time intervals. Colorimetric MTT assay was performed according to (Hansen J, Bross P, 2010). The absorbance of color in the solution was analyzed at 540 nm with micro-plate reader machine to calculate the 50% cytotoxic concentration (CC50), effective concentration (EC50) and viability of the cells.

#### 3.7.3 Percentage of protection

After one hour of incubation with EC50 of test compounds in different types of exposure with 100 TCID50 of the virus, the cells were washed with 1X PBS and, then medium supplemented with Trypsin_TPCK was added (100μl/well). After 48 hours of incubation at 37°C, viabilities of the cells were evaluated by MTT. The percentage of protection of each compound was calculated, in Microsoft Office Excel 2010 and SPSS, from the MTT results of mock-infected and infected cells after 48 hours of exposure, using the following formula:

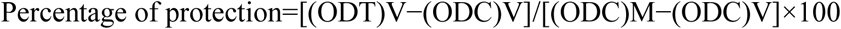

Where (ODT)V absorbance of test infected with virus, (ODC)V absorbance of negative control with virus and (ODC)M absorbance of positive control with only cells and medium imply the absorbance of the treated sample, the virus-infected control (no compound) and the negative control (no virus and no compound), respectively.

## Results

In this project of commercial micronutrient combination known as commercial iodine complex was evaluated for mutagenicity, cytotoxicity and anti-viral activity. The project also included the same parameters for the comparable micronutrient combination prepared in our laboratory. Moreover, individual ingredients were also tested for their antiviral, cytotoxic and mutagenic potential.

### 4.1 Cytotoxicity assays

The quantification of Chicken Embryo Fibroblasts was calculated by using hemocytometer and results are present in **Table 1**. It is important to note that the results for the next steps of cytotoxicity effects are shown in **Figure 1**.

The results for **Cytotoxic activity of MTI commercial iodine complex** are given in the **Table 2**. Ten concentrations of MTI commercial iodine complex, 187ug/ml, 93, 46.5, 23, 11.5, 5.8, 2.9, 1.4, 0.7 and 0.35 all units in ug/ml were tested for cytotoxicity and Cell Survival Percentage CSP was calculated. Respective CSP in percentage was 25.148*, 51.957, 59.311, 61.565, 64.175, 65.599, 78.054, 81.376, 83.867, 86.832 in percent values. Results clearly indicate that higher concentrations of 187ug/ml, showed cell survival percentage of less than 50% mean this concentration is higher than 50% cytotoxic concentrations and contain potential for cytotoxicity.

**Cytotoxic activity of UVAS Iodine complex** was determined at concentration 35.6µ g/ml, 17.78µ g/ml, 8.9µ g/ml, 4.5µ g/ml, 2.3µ g/ml, 1.1µ g/ml, 0.5µ g/ml, 0.2µ g/ml,0.13µ g/ml and 0.06 µ g/ml **(Table 3)**. Results clearly indicate that higher concentrations of 35.6ug/ml, 17.8ug/ml, 8.9ug/ml and 4.523ug/ml showed cell survival percentage of less than 50% mean these concentrations are higher than 50% cytotoxic concentrations (CC50) and contain potential for cytotoxicity.

**Cytotoxic activity of iodine** was determined at concentration 22.4 µ g/ml, 11.2µ g/ml, 5.6µ g/ml, 2.8µg/ml, 1.4µ g/ml, 0.7µ g/ml, 0.35µg/ml, 0.175µg/m,l 0.08µ g/mland 0.04µg/ml (**Table 4**). Results clearly indicate that higher concentrations of 22.4ug/ml, 11.2ug/ml and 5.6ug/ml showed cell survival percentage of less than 50% mean these concentrations are higher than 50% cytotoxic concentrations (CC50) and contain potential for cytotoxicity.

In the last three cases, while rest of the seven concentrations showed cytotoxicity less than 50% and have concentrations above CC50 and can be rendered quite safe concentrations.

**Cytotoxic activity of Ribavirin, Amantadine and Oseltamivir** were determined at concentration 100µ g/ml, 50µ g/ml, 25µ g/ml, 12.5µ g/ml, 6.2µ g/ml, 3.1µ g/ml, 1.6µ g/ml, 0.8µ g/ml, 0.4µ g/ml, and 0.2µ g/ml (**Table 5**). Results indicate that all the concentrations showed cytotoxicity less than 50% and Cell Survival Percentage was higher than 50% indicating that even at higher concentrations. Ribavirin, Amantadine and Oseltamivir are not cytotoxic and relatively safe values have been obtained in the experiment.

### 4.2 Anti-Viral Assay

Cell culture technique was performed to evaluate antiviral activity of commercial micronutrient combination Commercial iodine complex and laboratory made micronutrient combination UVAS-Commercial iodine complex along with their individual ingredients. Ribavirin, Amantadine and Oseltamivir were used as standards. Antiviral activity was measured against H9N1 influenza virus by using MTT Assay on Chicken Embryo Fibroblast cells. The antiviral activities comparison in terms of cell survival percentage are shown in **Figure 2**.

**Antiviral activity of MTI Commercial iodine complex** showed that concentrations 93µ g/ml,46.5µ g/ml, 23µ g/ml, 11.5 µ g/ml, 5.8µ g/ml, and 2.9µ g/ml had antiviral activity against H9N1 influenza virus as cell survival in these concentrations was above 50% while concentration 187 µ g/ml, had no antiviral activity as the cell survival was found out to be below 50 % and that might be due to cytotoxicity of micronutrient combination itself (**Table 2**). Concentrations of 1.4µ g/ml, 0.7 µ g/ml 0.35µ g/ml showed no antiviral activity as Cell Survival Percentage was much less than 50%.

**Anti-Viral activity of UVAS Commercial iodine complex** was determined at concentration 35.6µg/ml, 17.8µ g/ml, 8.89µ g/ml, 4.44µ g/ml, 2.22µ g/ml, 1.11µ g/ml, 0.55µ gm/ l, 0.27µ g/ml,0.13µ g/ml and 0.06µ g/ml (**Table 3)**. The results showed that concentrations 2.22 µ g/ml and 1.11µ g/ml had antiviral activity against H9N1 influenza virus as cell survival in these concentrations was above 50% while concentration.4.44µ g/ml and 0.55ug/ml also showed some activity as cell survival percentage was around 50%.

**Antiviral activity of iodine** was determined at concentration 22.4µ g/ml, 11.2µ g/ml, 5.6µ g/ml, 2.8µ g/ml, 1.4µg/ml, 0.7µ g/ml, 0.35µ g/ml, 0.175µg/ml, 0.08µ g/ml and 0.04µ g/ml (**Table 4**). 5.6µ g/ml and 0.175ug/ml also showed some activity as cell survival percentage was around 50 percent but on lower side while 22.4µ g/ml, 11.2µ g/ml,0.08µ g/ml and 0.04µ g/ml had no antiviral activity as the cell survival was found out to be below 50 % and that might be due to cytotoxicity of micronutrient combination itself or suboptimal concentrations for antiviral effect.

**Antiviral activity of Ribavirin, amantadine and oseltamivir** was determined at concentration 100µ g/ml, 50µg/ml, 25µ g/ml, 12.5µ g/ml, 6.2µ g/ml, 3.1µ g/ml, 1.6µ g/ml, 0.8µ g/ml,0.4µg/m and 0.2µ g/ml (**Table 5**). The results showed that concentrations 100µ g/ml, 50µ g/ml,25µ g/ml, 12.5µ g/ml, 6.25µ g/ml, 3.1µ g/ml and 1.6µ g/ml, had antiviral activity against H9N1 influenza virus as cell survival in these concentrations was above 50% while concentration.0.8µ g/ml, 0.4µ g/ml also showed some activity as cell survival percentage was around 50 percent but on lower side while 23.426ug/ml l had no antiviral activity as the cell survival was found out to be below 50 % and that might be due to suboptimal concentrations for antiviral effect. On the other hand, in amantadine, the values 1.6ug/ml, 0.8µ g/ml, 0.4µ g/ml and 0.2ug/ml had no antiviral activity as the cell survival was found out to be below 50 % and that might be due to suboptimal concentrations for antiviral effect. The results obtained with oseltamivir that concentrations 100µ g/ml, 50µ g/ml,25µ g/ml, 12.5µ g/ml, 6.25µ g/ml, 3.1µg/ml, 1.6µ g/ml, 0.8µg/ml, 0.4µg/ml and0.2µ g/ml all had antiviral activity against H9N1 influenza virus as cell survival in these concentrations was above 50% while concentration. This shows that even lowest concentrations of oseltamivir have the capability to inhibit influenza virus strain and possess highly active antiviral properties.

### 4.3 Mutagenicity assays

Mutagenicity potential of micronutrient combinations, individual ingredients and standard drugs was determined by **Ames** *Salmonella* microsomal assay in relation with the number of revertant colonies per plate. The Mutagenic Index of test chemicals was determined by following formula:

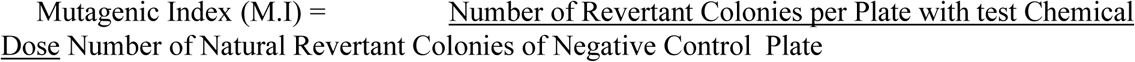

If its value is >2, it means the test concentration will be mutagenic. Mutagenic response was considered positive when number of colonies in test chemical plate were ≥ two-fold than the natural revertant of negative control or background colonies (Bajpayee et al. 2006).

The results for **mutagenic potential of MTI Commercial iodine complex**, **Mutagenic potential of UVAS-Iodine complex**, **Iodine+Potassium Iodide, Ascorbic Acid, Ribavirin, Amantadim and Oseltamivir** were as follows while using TA100 and TA 98 strains both with and without metabolic activation mixture S-9. They indicated clearly that no single dilution showed MI values equal to or greater than 2 hence all the concentrations can be rendered non-mutagenic in lieu of these results through Ames test although other methods may reveal different results.

## Discussion

A drug or combination being tested for antiviral activity should also be tested for cytotoxic activities to single out the concentrations of a chemical which produce antiviral effect yet do not produce cytotoxicity (Gqaleni et al. 2012).

**Ribavirin** is used as standard antiviral drug and an antiviral agent effective against a wide variety of viruses, including Hepatitis C Virus and influenza virus. It is effective in treatment and prevention of oseltamivir resistant influenza at 75mg/kg/day and 30mg/kg doses respectively (Smee et al. 2012). Cytotoxicity studies of ribavirin have been controversial in terms of high dose toxicity. Animal studies have demonstrated that 70mg/kg (roughly equivalent to 110ug/ml) or higher doses have been proven highly cytotoxic and genotoxic in bone marrow and spermatids (Narayana, D’Souza & Seetharama Rao, 2002). More studies have demonstrated cytotoxic effects of ribavirin on immune cells as part of its liver protective effect. Our studies demonstrated pronounced effect of ribavirin against H9 influenza virus. Although concentrations below 1ug/ml did not show significant antiviral activity but concentrations above 1ug/ml were effective against influenza virus which augment previous research about antiviral activity of ribavirin. The effect of ribavirin is mostly attributed to inhibition of inositol monophosphate dehydrogenase pathway thus blocking GTP conversion and consequently production of viral proteins. However, in vivo efficacy can be more profound due to immune system activation or other coordinating mechanisms (Krajczyk et al., 2013).

Another ingredient studied for cytotoxicity was **iodine**. Iodine have long been used as antiseptic due to its ability to kill microbes like MRSA, *enterococcus, chlamydia* and most of viruses. Concentrations of 2.8ug/ml, 1.4ug/ml, 0.7ug/ml and 0/35ug/ml showed prominent antiviral activity, showed no mutagenic potential and were not cytotoxic. It has shown viricidal activity against herpes simplex, adeno- and enteroviruses, HIV, HCV and H5N1 influenza virus (Sabracos et al. 2007). Most of the studies regarding iodine has been done with povidone-iodine, which is commonest form applied in clinics as antiseptics. Although 1000-fold dilution, 7.7ug/ml did not show toxicity but concluded that given more exposure, the lower concentration could be toxic (Schmidlin et al. 2009). Present study indicated that higher concentrations may be cytotoxic for the cells harboring virus and may not show cell survival which might have been there due to antiviral activity. Our results involving iodine alone were not much of encouragement as iodine + potassium iodide combination demonstrated toxicity at higher concentrations.

**MTI Commercial iodine complex** is the major test drug which had different properties at different levels. It showed cytotoxicity at highest concentrations of 187ug/ml while the same concentration was non-mutagenic and did not show any antiviral activity. Commercial iodine complex, a micronutrient combination containing iodine, potassium iodide, NaCl and Ascorbic acid along with excipients as its active constituents claim to possess antiviral activity. Its active ingredients have demonstrated weak antiviral activities in various studies (Shatzer et al. 2013). Specifically, ascorbic acid has been implied in antiviral activities against many viruses’ in vitro and in vivo studies. A study concluded that it possesses excellent efficacy against influenza virus too (Jariwalla et al., 2007). In this context, it should be no surprise if Commercial iodine complex containing ascorbic acid and high levels of iodine show significant antiviral activity.

UVAS-Iodine complex concentrations (the combination prepared in our laboratory that simulated the composition of commercial preparation) showed cell survival percentage more than 50 percent. In stark contrast, commercial micronutrient combination, Commercial iodine complex showed very less toxicity comparatively. It showed no traces of cytotoxicity at lower concentrations and although cell survival percentage decreased with the increase in concentration, it never reached to CC50 until last and highest concentration. In comparison, simple iodine + potassium iodide combination and laboratory prepared combination showed lower values for only up to 6 dilutions while commercial preparation showed no cytotoxicity up to 9 dilutions and only 10thth highest dilution made it to the cytotoxic ranks. It becomes more perplexing by the fact that all three combinations have similar amount of iodine, a major source of cytotoxicity, so how results could be different. One plausible explanation may be presence of ascorbic acid in commercial preparation which is shown to decrease the cytotoxicity of many otherwise cytotoxic compounds (Das Roy et al. 2013; Park et al. 2012). But in this case, UVAS-Iodine complex should also show similar effect which it does not. Another explanation might be the origin of iodine. If iodine is obtained from seaweeds or other biological sources, it is believed to be less cytotoxic as demonstrated in animal studies and case studies involving inhabitants of Japanese islands consuming large amount of iodine rich seaweeds.

The findings may be supported by presence of an additional undisclosed antiviral ingredient in excipients of commercial preparation as part of pharmaceutical company’s policy. Another likelihood is non-mineral source of iodine. In various studies, it has been demonstrated that bioactive iodinated compounds found in living organisms (Misurcova et al. 2011).

Production of genetic aberrations is the consequence of interaction of many chemicals with cells. Genetic aberrations can be measured by many tests including bone marrow analysis, rodent-micronucleus formation, comet assay and bacterial mutation reversion assay. *E. coli* and *Salmonella typhimurium* genetic mutation reversion assay or Ames test has been robust at detecting potential mutagenic chemicals which can subsequently cause carcinogenicity (Zeiger, 2013).

Ribavirin is proven to be mutagenic in different studies while utilizing different techniques. It has caused mutagenicity in rodent bone marrow and has produced micronucleus. Recent studies have concluded that its main antiviral effect is due to production of mutations in viral genome rather than inhibition f IMPDH (Crotty et al. 2002). But it has been noted that ribavirin has never shown significant reverse mutation through *Salmonella*/Ames assay even up to concentrations of 200ug/ml and Ames test has declared ribavirin as a non-mutagenic chemical (PEGASYS literature from Roche). Our studies confirm the finding that significant mutagenic potential was not observed while studying it in our lab using TA98 and TA100 strains of *Salmonella* in the presence and absence of liver extract preparations of S9.

In this project mutagenicity of COMMERCIAL IODINE COMPLEX (micronutrient combination) a micronutrient combination, UVAS-Iodine complex and individual ingredients including ascorbic acid and iodine were tested against TA 100 and TA 98 strains of *Salmonella typhimurium*, no significant number of revertant colonies were present when testing Commercial iodine complex, UVAS-Iodine complex and individual elements against TA 100 and TA 98 as compared to negative control plate with and without using metabolic activation system S-9. It signifies that nutrients caused only base pair substitution mutation at in number of colonies equivalent to that of negative control or bit higher but not statistically or experimentally significant quantities of revertant colonies were produced. Although the number of colonies in TA100 with and without S9 metabolic activation system were higher when compared to case of TA 98, significant number of revertant did not appeared in comparison with the negative control plate. Literature review has suggested that these micronutrients are not found to be mutagenic in bacterial Ames test. Iodine as an ingredient might show genotoxicity or other DNA defects at much higher doses than used in this experiment but still has not found to be mutagenic in bacterial Ames assay (Dusman et al. 2011). Reason for iodine toxicity lies in the fact that it could produce free radicals which may cause membrane damage and DNA breaks. The repairing system may not be able to cope with the extent of mutations and could leave some base pairs unrepaired (Fischer et al. 2011). On the opposite side, ascorbic acid is not only found to be non-mutagenic by Ames and other assays but also is found to decrease mutagenic potential of many powerful mutagens when combined together. Quite a notable number of studies reveal that concomitant administration of vitamin C could reduce DNA mutations and DNA damage caused by potential genotoxic, mutagenic substances (Das Roy et al., 2013; Park et al., 2012). Although least likely, but micronutrient combination may have mutagenic potential which is successfully reversed due to presence of ascorbic acid. Ironically, very few but recent studies have demonstrated ascorbic acid to be an important cause of producing mutations and DNA damage while this potential is obtained only when high pro-oxidant concentrations are achieved and is limited only to cancer and virus infected cells. Further advancements are soon to be expected in this regard (Mamede et al. 2012).

On the other hand, standard drugs used along with these micronutrient combinations did not show significant mutagenicity. Ribavirin, Amantadine and Oseltamivir were tested along with our test preparations produced revertant bacterial colonies within range to negative control. Amantadine have been shown to be strictly non-mutagenic even at highest concentrations possible in human body. Our results support that situation. Same is true for oseltamivir which is not mutagenic at therapeutic and higher concentrations (Ila & Ilhan, 2012). Ribavirin has been tested widely for its mutagenicity through different models. Interestingly, it has proven to be a strong mutagenic drug when tested in-vivo animal models at concentrations not much higher than therapeutic concentrations, but it has always failed to produce positive results in bacterial Ames test even at 5000 times higher concentrations than therapeutic ones (Krajczyk et al., 2013).

According to the t-test applied for statistical analysis mutagenic potential showed no significant difference in case of TA 98 with and without S-9 mix as p value was less than 0.05. Whereas no significant difference was also there in case of TA 100 with and without S-9 mix. Overall Base pair substitution mutation values were high as compared to Frame shift mutation.

When T-test was applied for the statistical analysis of data, comparison revealed that there was no significant difference (p > 0.05) between the mutagenic potential of combination doses as compared to mutagenic potential of nutrients alone. In case of comparison between combination and iodine the difference was slightly significant only when mutagenic potential was compared against TA 100 without S-9.

## Conclusions

Micronutrients, essential nutrients to organism manutention that are needed in small amounts, are as important for life as macronutrients. In this study, we tested cytotoxicity, antiviral effect and mutagenicity of different compounds.

Ribavirin’s highest six concentrations ranging from 1.6ug/ml to 100ug/ml shoed non-mutagenicity, antiviral efficacy and were not cytotoxic. It showed no cytotoxicity at all concentrations. MTI Commercial iodine complex is the major test drug which had different properties at different levels. It showed cytotoxicity at highest concentrations of 187ug/ml while the same concentration was non-mutagenic and did not show any antiviral activity. UVAS Commercial iodine complex was the laboratory made preparation which simulated the composition of commercial preparation and the effects depends of the exactly concentration.

To our knowledge, the combination consisting of iodine, ascorbic acid, potassium iodide is effective in the treatment of viral infections. Our findings highlight that nationally representative data are needed to control the development of nutrition and public health programs, such as dietary diversification, micronutrient fortification and supplementation.

## Acknowledgements

We are really thankful for department of pharmacology, and research laboratories of university of veterinary and animal sciences to let us use facilities. We are also thankful for MTI medical to provide funding for the execution of this research project.

---

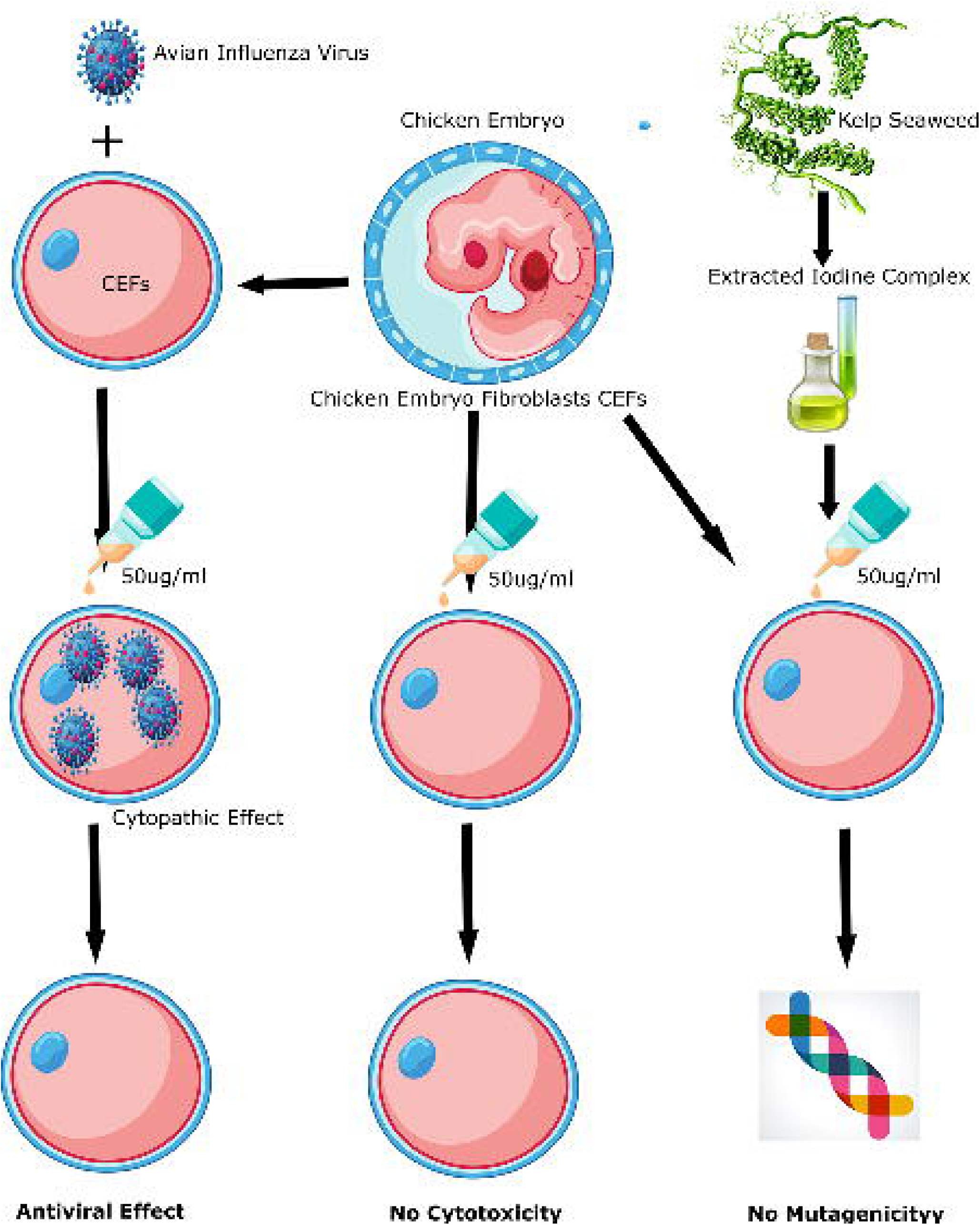

