## Supplemental Tables and Figures for "Evaluation of cytotoxic, antiviral effect and mutagenic potential of a micronutrient combination in vitro cell culture"

**Table 1** - Cytometry of Chicken Embryo Fibroblasts at the time of Assay.

|  | Number of live cells | Number of dead cells | Total |
| --- | --- | --- | --- |
| Square 1 | 9 | 3 | 12 |
| Square 2 | 9 | 5 | 14 |
| Square 3 | 16 | 2 | 18 |
| Square 4 | 11 | 1 | 12 |
| Total | 50 | 11 | 61 |

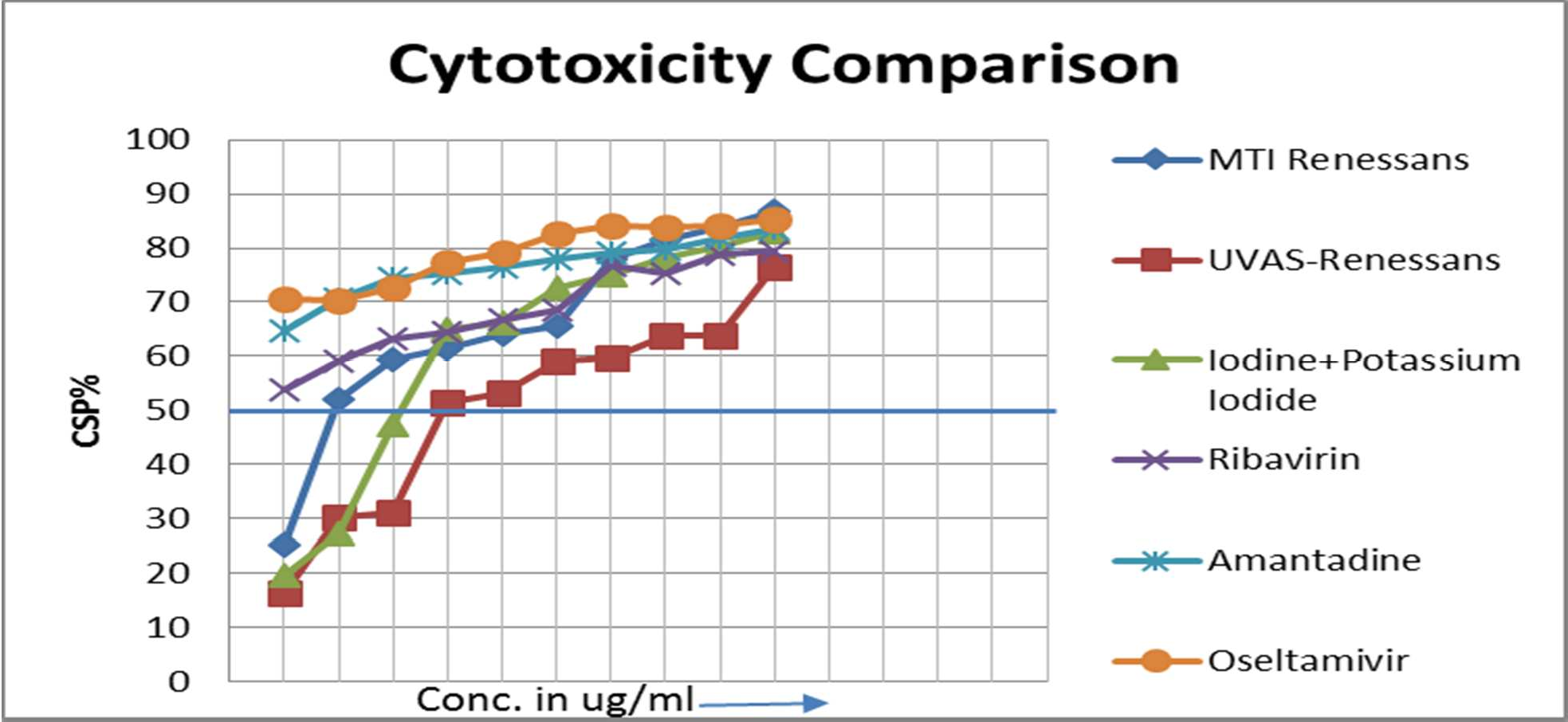

**Figure 1** - Comparison of cytotoxicities of MTI Renessans, UVAS-Renessans, Ribavirin, Amantadine and Oseltamivir analyzed in this study.

**Table 2** - Cytotoxicity, Antiviral and Mutagenicity profile comparison for Commercial Iodine Complex MTI Renessans

| Commercial Iodine Complex Renessans | CSP  Cytotoxicity | CSP  antiviral | Mutagenicity | Comments |
| --- | --- | --- | --- | --- |
| 187 | 25.148* | 15.776 | Non-Mutagenic | Cytotoxic, No antiviral effect |
| 93 | **51.957** | **73.546** | **Non-Mutagenic** | Non-cytotoxic, Antiviral |
| 46.5 | **59.311** | **80.545** | **Non-Mutagenic** | Non-cytotoxic, Antiviral |
| 23 | **61.565** | **74.851** | **Non-Mutagenic** | Non-cytotoxic, Antiviral |
| 11.5 | **64.175** | **53.736** | **Non-Mutagenic** | Non-cytotoxic, Antiviral |
| 5.8 | **65.599** | **54.329** | **Non-Mutagenic** | Non-cytotoxic, Antiviral |
| 2.9 | **78.054** | **54.685** | **Non-Mutagenic** | Non-cytotoxic, Antiviral |
| 1.4 | 81.376 | 12.692 | Non-Mutagenic | Non-cytotoxic, Low antiviral effect |
| 0.7 | 83.867 | 15.658 | Non-Mutagenic | Non-cytotoxic, No antiviral effect |
| 0.35 | 86.832 | 13.285 | Non-Mutagenic | Non-cytotoxic, No antiviral effect |

**Table 3** - Cytotoxicity, Antiviral and Mutagenicity profile comparison for UVAS-iodine complex

| UVAS  Iodine complex | CSP  cytotoxicity | CSP  Antiviral | Mutagenicity | Comments |
| --- | --- | --- | --- | --- |
| 35.56 | 16.251 | 6.524 | Non-Mutagenic | Cytotoxic, No antiviral effect |
| 17.78 | 30.249 | 16.846 | Non-Mutagenic | Cytotoxic, No antiviral effect |
| 8.89 | 31.079 | 44.483 | Non-Mutagenic | Cytotoxic, No antiviral effect |
| 4.44 | **51.483** | **55.516** | **Non-Mutagenic** | Non-cytotoxic, Antiviral |
| 2.22 | **53.143** | **58.956** | **Non-Mutagenic** | Non-cytotoxic, Antiviral |
| 1.11 | **58.956** | **63.226** | **Non-Mutagenic** | Non-cytotoxic, Antiviral |
| 0.55 | 59.667 | 37.722 | Non-Mutagenic | Non-cytotoxic, Very little antiviral  effect |
| 0.27 | 63.819 | 18.742 | Non-Mutagenic | Non-cytotoxic, Non-antiviral effect |
| 0.13 | 63.819 | 5.3380 | Non-Mutagenic | Non-cytotoxic, Non-antiviral effect |
| 0.06 | 76.512 | 7.591 | Non-Mutagenic | Non-cytotoxic, Non-antiviral effect |

**Table 4** - Cytotoxicity, Antiviral and Mutagenicity profile comparison for Iodine+Potassium Iodide.

| Iodine+  Potassium  Iodide | CSP  toxicity | CSP antiviral | Mutagenicity | Comments |
| --- | --- | --- | --- | --- |
| 22.4 | 19.545 | 17.207 | Non-Mutagenic | Cytotoxic, No antiviral effect |
| 11.2 | 27.284 | 12.574 | Non-Mutagenic | Cytotoxic, No antiviral effect |
| 5.6 | 47.331 | 39.739 | Non-Mutagenic | Cytotoxic, No antiviral effect |
| **2.8** | 65.005 | 50.102 | Non-Mutagenic | Non-cytotoxic and almost IC50  antiviral |
| **1.4** | 66.073 | 51.838 | Non-mutagenic | Non-cytotoxic and almost IC50 antiviral |
| **0.7** | 72.597 | 69.513 | Non-Mutagenic | Non-cytotoxic and antiviral effect |
| **0.35** | 75.088 | 68.683 | Non- Mutagenic | Non-cytotoxic and antiviral effect |
| 0.175 | 78.291 | 40.688 | Non-Mutagenic | Non-cytotoxic and no antiviral  effect |
| 0.08 | 80.189 | 36.061 | Non-Mutagenic | Non-cytotoxic and no antiviral  effect |
| 0.04 | 83.036 | 12.692 | Non-Mutagenic | Non-cytotoxic and no antiviral  effect |

**Table 5** - Cytotoxicity, Antiviral and Mutagenicity profile comparison for Ribavirin

| **Ribavirin** | **CSP**  **Cytotoxicity** | **CSP Antiviral** | **Mutagenicity** | **Comments** |
| --- | --- | --- | --- | --- |
| 100 | 53.879 | 57.739 | Non-Mutagenic | Non-cytotoxic, Antiviral |
| 50 | 59.047 | 77.010 | Non-Mutagenic | Non-cytotoxic, Antiviral |
| 25 | 63.262 | 79.356 | Non-Mutagenic | Non-cytotoxic, Antiviral |
| 12.5 | 64.329 | 88.398 | Non-Mutagenic | Non-cytotoxic, Antiviral |
| 6.2 | 66.785 | 87.511 | Non- Mutagenic | Non-cytotoxic, Antiviral |
| 3.1 | 68.446 | 83.900 | Non- Mutagenic | Non-cytotoxic, Antiviral |
| 1.6 | 76.683 | 86.478 | Non-Mutagenic | Non-cytotoxic, Antiviral |
| 0.8 | 75.444 | 47.686 | Non-Mutagenic | Non-cytotoxic, Low antiviral effect |
| 0.4 | 78.766 | 46.062 | Non-Mutagenic | Non-cytotoxic, Low antiviral effect |
| 0.2 | 79.478 | 23.426 | Non-Mutagenic | Non-cytotoxic, No Antiviral effect |

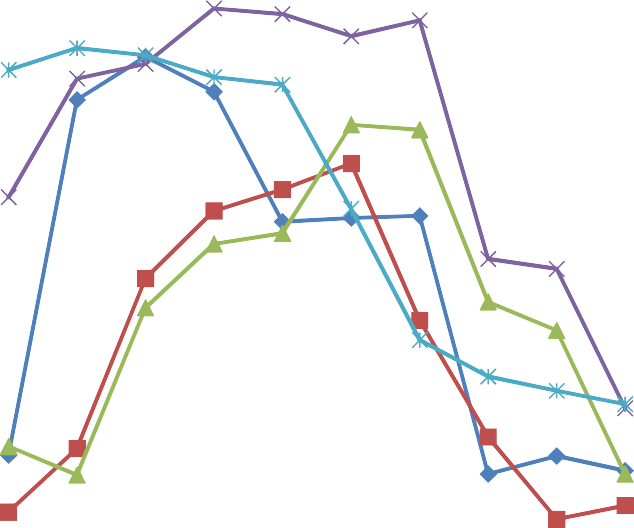

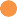

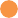

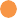

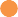

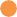

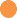

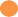

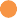

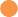

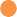

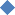

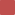

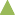

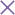

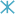

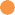

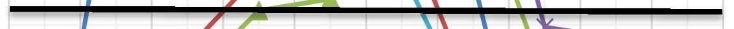

**ANTIVIRAL COMPARISON**

100

90

RENESSANS

80

70

UVAS_Renessans

60

Iodine plus Potassium Iodide

50

40

Ribavirin

30

Amantadine

20

10

Oseltamivir

0

1

2

3

4

5

6

7

8

9	1 0

CELL SURVIVAL PERCENTAGE CSP%

**Figure 2** - Antiviral activities comparison in terms of cell survival percentage between MTI Renessans, UVAS Renessans, Ribavirin, Amantadine and Oseltamivir.
